# Parental care enhances reproductive success without remodeling the carcass microbiome in a wild burying beetle

**DOI:** 10.64898/2026.07.16.739041

**Authors:** Wei-Jiun Lin, Chi-Heng Hsieh, Syuan-Jyun Sun

## Abstract

Nutrient cycling depends on interactions between insect decomposers and the microbial communities that colonize ephemeral resources such as animal carcasses. Burying beetles (*Nicrophorus* spp.) are thought to manipulate carcass microbiomes to facilitate reproduction, yet most experimental work relies on a few standardized laboratory carcasses that obscure the ecological heterogeneity beetles face in the wild. Using 76 carcasses from 27 vertebrate species of birds, mammals, and reptiles, we manipulated *Nicrophorus nepalensis* parental care and characterized carcass, adult-gut, and larval-gut microbiomes by 16S rRNA sequencing. Parental care strongly increased breeding success and brood mass, and buffered reproductive success against variation in carcass identity. By contrast, carcass type and decomposition stage, not care, primarily structured the carcass microbiome; a modest care signature emerged only late in decomposition, was strongest in the adult gut, and was absent from the larval gut. Parental care thus acts principally as a direct driver of reproductive success rather than by remodeling the carcass or larval microbiomes.

## Introduction

Nutrient cycling in terrestrial ecosystems largely relies on decomposition, with microbial communities playing a central role not only in the breakdown and transformation of organic matter, but also in shaping the structure and function of decomposition networks (Bardgett and Van Der Putten 2014). Animal carcasses represent nutrient-rich yet ephemeral and spatially unpredictable resource hotspots. Their decomposition releases pulses of nutrients and moisture into surrounding soil (Barton et al 2013), markedly altering local biochemical conditions and driving rapid microbial proliferation, community succession, and competitive interactions (Carter et al 2007, Cobaugh et al 2015, Metcalf et al 2016, Burcham et al 2024, von Hoermann et al 2022). Microbial assemblages on field-exposed carcasses are therefore highly heterogeneous (Barton et al 2013, Metcalf et al 2016). Scavenger and decomposer activity can further restructure carcass microbiota. For example, in burying beetles carcass preparation and parental care are predicted to stabilize or redirect microbial communities (Korner et al 2023).

Interactions between microbes and insect decomposers include both competitive and facilitative relationships (Holt et al 2024, Rombaut et al 2023). Yet, how carcasses, microbial communities, and insect decomposers interact remains poorly understood (Benbow et al 2025). Advances in sequencing technologies have revealed the remarkable diversity and complexity of animal-associated microbiota and demonstrated their influence on host behavior, development, immunity, and metabolism (Ezenwa et al 2012, McFall-Ngai et al 2013). In insects, long-established symbiotic associations have enabled the exploitation of nutritionally or chemically challenging resources, facilitating the colonization of novel ecological niches (Behar et al 2008, Currie et al 1999). For example, fungal symbioses in gall-inducing insects have facilitated host-plant expansion and diversification (Joy 2013). However, it remains largely unclear how microbial communities on ephemeral resources, such as carcasses, feedback to shape the microbiota and fitness, especially across life stages in decomposer insects (Metcalf et al 2016, Wong et al 2015).

Burying beetles (*Nicrophorus* spp.) provide a tractable model system to investigate these interactions because they breed on small vertebrate carcasses and actively manage the resource. Adults remove fur or feathers, shape the carcass into a ball, apply oral and anal secretions, and bury it underground, which together help preserve carcass quality and enhance larval survival (Arce et al 2013, Cotter et al 2011, Pukowski 1933). Laboratory studies have demonstrated that burying beetles actively manipulate carcass microbial communities through antimicrobial secretions and microbial seeding, which can preserve resources and facilitate larval development (Rozen et al 2008, Duarte et al 2018, Shukla et al 2018a, Vogel et al 2017, Kaltenpoth and Steiger 2014).

Parents also use the carcass as a medium to transmit a core microbiota to their offspring (Shukla et al 2018b), a process that includes direct transmission of maternal gut bacteria to larvae during care, with larvae that do not receive care instead acquiring bacteria from the carcass itself (Wang and Rozen 2017), and parental care plays a crucial role in stabilizing larval gut communities (Ayayee et al 2025, Miller et al 2021). While these actions are known to preserve the resource, their precise influence on the temporal dynamics of the carcass microbiome remains unclear.

Most experimental work on burying beetles and their carcass microbiomes has relied on a small number of standardized laboratory carcasses, typically frozen mice or day-old chicks (Shukla et al., 2018a, 2018b; Wang and Rozen, 2017). However, different vertebrate carcasses vary substantially in nutrient composition, immune molecules, skin or feather chemistry, and resident microbial communities. Because these properties can shape both the microbiome available to beetles and the resource quality underlying breeding success, findings from a single laboratory carcass type may not generalize across the taxonomically diverse carcasses beetles encounter in the wild.

Furthermore, field carcass communities can be highly complex and variable. A previous analysis of an overlapping wild-carcass panel found that carcass size, rather than source or taxonomic identity, predicted breeding performance (Hsu et al 2024); however, that analysis did not test whether this relationship depends on the presence of parental care. This distinction is important because parental care may buffer or amplify resource-associated effects on offspring production (Metcalf et al 2016, Pechal et al 2013).

Here, we investigate how parental care, carcass identity, and decomposition stage shape the carcass microbiome, the gut microbiomes of adults and larvae, and reproductive success. To do so, we conducted a breeding experiment manipulating the presence of parental care across 76 wild and laboratory-sourced carcasses spanning 27 species of birds, mammals, and reptiles, and characterized the resulting microbial communities via 16S rRNA gene sequencing at both early (Day 5) and late (Day 9) stages of decomposition, in addition to the adult and larval gut, before assessing breeding success and reproductive output. Specifically, we ask whether parental care, carcass identity, and their interaction predict both the microbiome and reproductive success, treating these as two related but distinct response systems. Whether parental care influences reproductive success primarily by modifying carcass microbial environments, transmitting beneficial microbes, or both remains unresolved. Specifically, we test (1) whether parental care and carcass type predict breeding success and reproductive output, including whether any effect of carcass type on success itself depends on the presence of parental care; (2) whether carcass type, decomposition stage, and parental care structure microbial community composition and diversity on the carcass, and in the adult and larval gut; and (3) whether carcass, adult-gut, and larval-gut microbial community structure are associated across compartments (carcass to adult gut, carcass to larval gut) and across time (Day 5 to Day 9). By sampling across a substantially wider host-taxonomic range, this study extends the taxonomic scope of prior work on parental care and the carcass microbiome in this genus.

## Materials and Methods

### Beetle rearing and breeding trials

The burying beetles (*Nicrophorus nepalensis*) used in this study were the descendants of wild-caught individuals collected in Taipei and New Taipei City, Taiwan, in 2023 and 2025. Beetles were reared in growth chambers maintained at 70% relative humidity on a 10:14 h light:dark cycle. Chamber temperature fluctuated between 16-20 degrees C (mean 17.8 degrees C) to mimic natural diurnal conditions during the field breeding season (November-April) in northern Taiwan. Each year during the field season, newly collected beetles were introduced into the existing laboratory colonies to limit genetic bottlenecks.

For breeding trials, unrelated male and female beetles were paired and placed in plastic containers (10.8 x 7.5 x 2.1 cm) half-filled with moist commercial potting mix. After 24 h of mating, females were transferred to transparent breeding boxes (14.2 x 6.3 cm) containing 2 cm of moist soil, with a defrosted carcass placed on the soil surface on Day 1. A total of 27 carcass species (76 individual carcasses, organized as 38 sibling-paired units) were used: two laboratory-sourced species (commercially purchased frozen mice and day-old chicks, n = 4 carcasses) and 25 wild-caught species obtained via a citizen-science network and the Wild Bird Society of Taipei, comprising 13 bird, 10 reptile, and 2 mammal species (body mass 3.1-78.4 g). Carcasses were collected and sequenced in two rounds: an initial round (batch 1, 40 carcasses) and a subsequent round (batch 2) that added 7 species (18 sibling-paired units) to increase sample size for comparisons. Each carcass species was used in both the parental care and no parental care treatments. To minimize parental family effects, a sibship-pairing scheme (e.g., males from family X with females from family Y) was used across treatments (Sun et al 2020). Wild carcasses had known accidental causes of mortality (e.g., traffic or window collision) within the preceding four months and were not associated with poisoning or illness. In paired treatments, both carcasses were required to be of the same species, with body mass differing by <=20% and collection sites within 3 km, to reduce among-pair variation in microbial communities. All breeding containers were maintained under identical environmental conditions in growth chambers.

In the no parental care treatment, females were removed on Day 5, at which point *N*. *nepalensis* typically completes oviposition and carcass preparation. At Day 12, trials were considered complete when larvae reached the third instar and at least one individual had exited the nest cavity, after which females and all surviving larvae were collected. Brood size was recorded as the number of surviving larvae, and brood mass was measured to the nearest 0.0001 g using an analytical balance (ATX224R, Shimadzu, Japan).

### Microbe sampling

Carcass sampling. Microbe samples were collected using sterile cotton swabs (FLOQSwabs Flocked Swabs 503CS01). Each swabbing lasted 10 s with repeated strokes, after which the swab tip was placed into a 1-ml phosphate-buffered saline (PBS) tube. On Days 5 and 9, swabs were taken from the feeding cavity; larvae were gently and temporarily displaced during sampling to minimize disturbance. All samples were stored at -20 degrees C until processing.

Beetle gut sampling. The adult female and one randomly chosen third-instar larva per brood were dissected immediately after collection using sterilized instruments, and the entire gut was removed, expressed onto a sterilized glass slide, and transferred into a 1-ml PBS tube. All samples were stored at - 20 degrees C until processing.

### Microbiome sequencing and bioinformatics

DNA was extracted from each swab sample and the V3-V4 region of the bacterial 16S rRNA gene was amplified and sequenced on an Illumina MiSeq platform (300 bp paired-end) by Genomics BioSci & Tech Co. (New Taipei City, Taiwan). Raw reads were adapter/primer-trimmed (Trimmomatic v0.39, Cutadapt v3.5) and quality-filtered; denoising, chimera removal, and amplicon sequence variant (ASV) inference were performed with DADA2 in QIIME2, and taxonomy was assigned with a SILVA v138 (99%) Naive Bayes classifier trained on the amplified V3-V4 region. Samples from the second collection round (batch 2) were processed with the same wet-lab and bioinformatic pipeline; ASV tables and taxonomy from the two batches were merged by ASV sequence identity, missing counts introduced by the merge were treated as true zeros, and any ASV lacking a Domain-level taxonomic assignment was excluded.

### Statistical analyses

All data processing and statistical analyses were conducted in R (version 4.4.2; R Core Team 2024).

### Sample classification

Each carcass was assigned to one of three taxonomic categories (bird, mammal, reptile) using an explicit, predefined species-to-category classification list (27 species: 14 bird, 3 mammal, 10 reptile). Each carcass was additionally classified by origin (wild-caught, n = 72, versus laboratory-sourced, n = 4) to evaluate carcass origin as a potential confound (Supplementary Results). A sequencing-batch variable distinguished the two collection/sequencing rounds and was included as a covariate in every community-level model.

### Diversity metrics

For each sample subset (carcass Day 5, carcass Day 9, adult beetle gut, larval gut), count tables were rarefied once to a fixed depth per subset (62,000-72,000 reads, set to the minimum sequencing depth observed within that subset) using vegan::rrarefy() (seed = 123; vegan package, Oksanen et al.). Alpha diversity was summarized as the Shannon index and observed ASV richness. Rarefaction curves and species accumulation curves (rarecurve(), specaccum()) were generated for each subset to confirm that sequencing depth was sufficient to approach saturation of richness (Fig. S6).

### Beta diversity and PERMANOVA

Bray-Curtis dissimilarities (vegdist()) were calculated on rarefied counts and ordinated by principal coordinates analysis (cmdscale()). Community composition was tested against carcass type, treatment (parental care vs. no care), decomposition stage, and sequencing batch using PERMANOVA (adonis2(), 999 permutations, marginal term testing), fitted separately for each sample subset (Day 5 carcass, Day 9 carcass, adult beetle gut, larval gut), with model terms varying by subset depending on which factors applied (full model specifications in Supplementary Methods). Pairwise carcass-type and treatment contrasts used the same approach with false-discovery-rate (Benjamini-Hochberg) correction across comparisons. Homogeneity of multivariate dispersion was checked for every grouping factor, including batch, with betadisper() and a permutation test (999 permutations), and carcass-type x treatment interaction terms were tested where sample size allowed.

A treatment x success interaction model, controlling for batch, was additionally fitted to evaluate whether microbial differences associated with treatment depended on breeding outcome.

### Alpha diversity models

Shannon diversity and observed richness were modeled as a function of carcass type, treatment, and batch. For subsets containing repeated sampling within the same nest of origin (Day 9, larvae), a linear mixed-effects model with nest as a random intercept (lme4::lmer(); lmerTest for approximate F/p-values; Bates et al. 2015) was fitted; where no nest contributed more than one sample, the model was automatically reduced to an ordinary linear model, because a random effect cannot be estimated without within-group replication. The Day 5 subset used ordinary linear models throughout, as no nest was resampled at that stage. Residual normality for every linear or mixed model was checked with the Shapiro-Wilk test, and results were interpreted alongside the corresponding non-parametric (Wilcoxon) tests when residuals departed from normality.

### Differential abundance

Differential ASV abundance between carcass types (Day 5) and between treatments (Day 9 and adult beetle gut) was tested with DESeq2 (Wald test on a negative-binomial GLM), retaining ASVs with a Benjamini-Hochberg adjusted p < 0.05. Differential abundance patterns were additionally summarized at the genus level to evaluate consistency among ASVs assigned to the same genus.

### Indicator species analysis

Indicator taxa for carcass type, treatment, and breeding success were identified using the IndVal.g statistic (indicspecies::multipatt(), 999 permutations) on rarefied counts.

### Cross-compartment and temporal comparisons

Mantel tests (999 permutations, Pearson correlation between Bray-Curtis distance matrices, matched by nest of origin) were used to test the association between carcass, adult-gut, and larval-gut community structure, and between Day 5 and Day 9 carcass communities (overall and within each treatment). Because carcass and adult-gut samples were often processed within the same sequencing batch, cross-compartment comparisons (carcass-adult gut, carcass-larval gut, adult gut-larval gut) were tested with partial Mantel tests controlling for batch (vegan::mantel.partial()), since unconditioned tests could not distinguish shared batch structure from true biological association (Supplementary Methods). For the adult gut-larval gut comparison, which remained significant after conditioning on batch alone, we additionally tested robustness to treatment-group structure with a partial Mantel test conditioning on treatment and a leave-one-out sensitivity analysis (Supplementary Methods). Within-nest microbiome turnover from Day 5 to Day 9 was quantified as the paired Bray-Curtis distance between the two time points and compared between treatments with a t-test, a Wilcoxon rank-sum test, and a linear model adjusting for carcass type and batch.

### Reproductive outcomes

Breeding success was defined as at least one surviving larva (brood size > 0) and analyzed with Fisher’s exact tests (success x treatment; success x carcass type) and binomial logistic regression (treatment, carcass type, and batch as predictors). Continuous reproductive measures (clutch size, brood size, brood mass, mean larval mass) were compared between paired treatment carcasses of the same species with Wilcoxon signed-rank tests, and, to allow adjustment for covariates, with linear models including treatment, carcass type, batch, and carcass mass.

To mirror a family-structured design, breeding success, brood size, and brood mass were additionally modeled with generalized linear mixed models including male and female family identity as crossed random intercepts ((1|male) + (1|female): binomial for success, Poisson for brood size, Gaussian (lmer) for brood mass). Brood mass was modeled once on the full dataset (including nests with zero surviving larvae, coded as brood mass = 0, for consistency with the brood-size and success models) and once restricted to successful nests only, so that whether treatment affects whether a brood succeeds at all is kept analytically separate from whether, given success, treatment affects how large the brood is.

To test whether the effect of carcass type on breeding success depended on treatment, a carcass x treatment interaction term was added to the binomial success model and compared with the corresponding main-effects-only model by a likelihood-ratio test; the same interaction was tested for brood size and brood mass. Carcass type was then examined separately within the No-care and Parent-care subsets using both standard and Firth’s penalized logistic regression, because several carcass-by-treatment cells were small enough to risk quasi-complete separation. Additional checks on this stratified comparison -- refitting on the original (batch 1 only) dataset and testing a nonlinear carcass-mass term -- are described in Supplementary Methods.

### Sensitivity and robustness analyses

To assess the robustness of the reproductive-outcome and community-composition results, models were repeated after excluding mammal carcasses, after excluding laboratory-sourced carcasses, with Firth’s penalized logistic regression, with the original (batch 1 only) dataset, and with a quadratic carcass-mass term. Full details of these analyses and their results are provided in Supplementary Methods and Supplementary Results.

## Results

### Parental care strongly increases breeding success and reproductive output

Breeding success was markedly higher under parental care (Fisher’s exact test, OR = 8.8, p < 0.0001). This effect remained highly significant after accounting for carcass type and sequencing batch (binomial GLM: OR = 16.1, 95% CI 4.6-77.9, p < 0.0001). Carcass type was also associated with success, with reptile and mammal carcasses more likely to succeed than bird carcasses (OR = 13.8 and 44.2, respectively; Table S4), although the mammal estimate was imprecise given the small mammal sample (n = 6 carcasses).

Brood size and brood mass were both higher under parental care once carcass type, batch, and (for brood mass) carcass mass were accounted for (linear models: brood size F = 15.6, df = 1,70, p < 0.001; brood mass F = 19.8, df = 1,70, p < 0.001; Fig. 1; Table S4). This effect on brood mass was no longer detected when the model was restricted to successful nests only (all p > 0.20), indicating that parental care primarily affects total reproductive output through increasing the likelihood of successful reproduction, rather than increasing offspring production among successful nests. Mean larval mass was unaffected by treatment (p = 0.29) but increased with carcass mass (p = 0.024).

**Figure 1.**
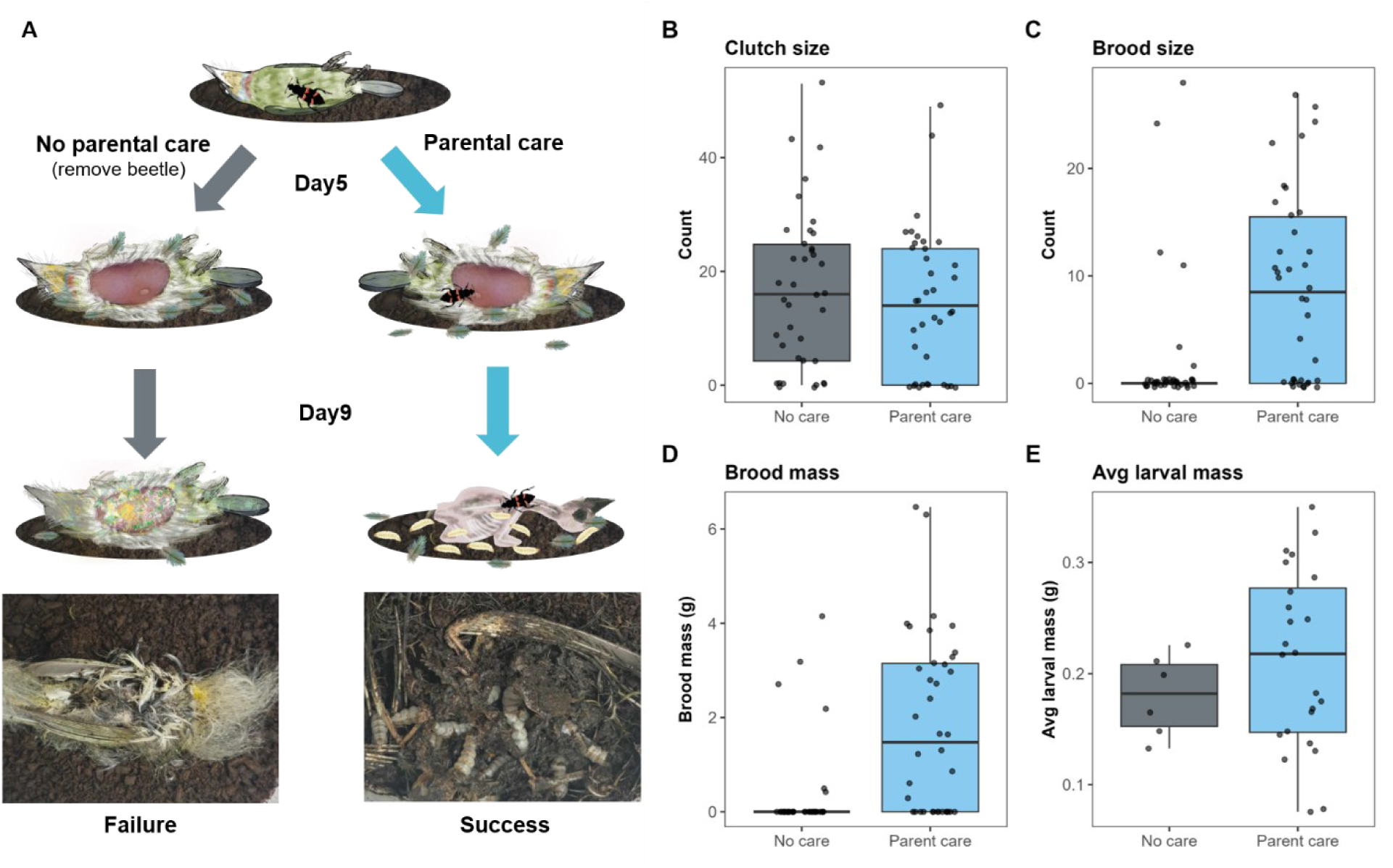
Effects of parental care on reproductive output. (A) Schematic of the experimental design: a carcass is assigned to the Parent-care or No-care treatment; in the No-care treatment the female is removed on Day 5. Representative nests are shown at Day 5 and Day 9, together with example successful and failed outcomes. Box plots of (B) clutch size, (C) brood size, (D) brood mass, and (E) mean larval mass for the No parental care and Parental care treatments.

The effect of carcass type on breeding success was not uniform across treatments. A formal carcass type x treatment interaction term was not statistically significant in the pooled logistic model (likelihood-ratio test, p = 0.827). However, stratifying by treatment showed that carcass type significantly predicted success within the No-care group (binomial GLM, OR = 26.0 for both mammal and reptile relative to bird, p = 0.042 each; Firth’s penalized regression likelihood-ratio test, p = 0.018) but not within the Parent-care group (Firth likelihood-ratio test, p = 0.248), with a similar pattern for brood mass (No-care: p = 0.044; Parent-care: p = 0.174). These stratified analyses provide exploratory evidence that carcass-type effects may be more pronounced in the absence of parental care, although this pattern was not supported by the formal interaction test.

### Community composition is structured primarily by carcass type and decomposition stage

Across the full carcass dataset (Day 5 and Day 9 combined), community composition was structured by carcass type (PERMANOVA, R^2^ = 0.056, p = 0.003) (Fig. 2) and decomposition stage (R^2^ = 0.093, p = 0.001) (Table S1); treatment alone was not significant (R^2^ = 0.018, p = 0.168), nor was a treatment x success interaction (R^2^ = 0.017, p = 0.346), and neither term reached significance within the Day 5 or Day 9 subsets considered separately (all p > 0.10).

**Figure 2.**
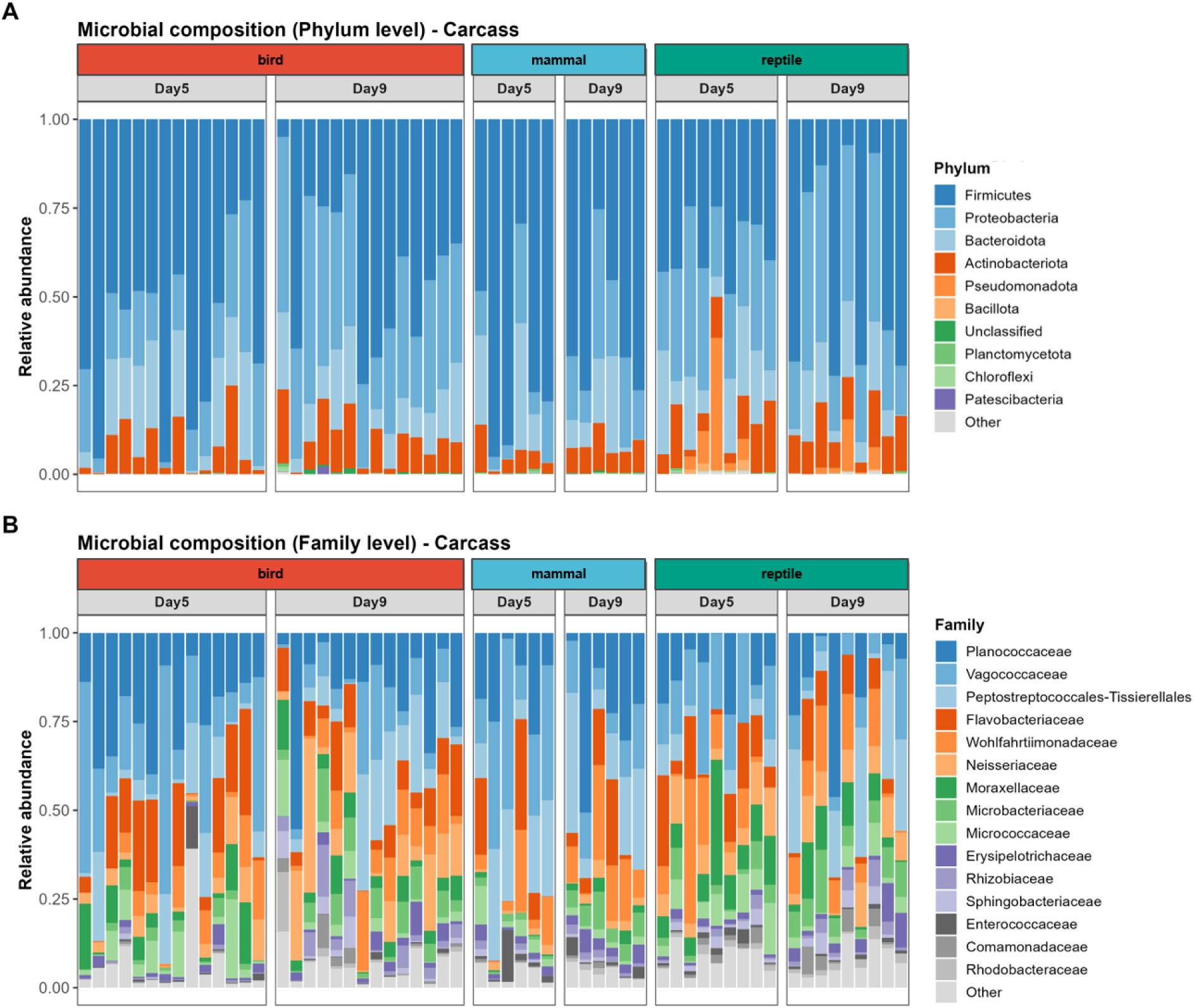
Carcass microbiome composition differs primarily by carcass type and decomposition stage. Relative abundance of (A) bacterial phyla and (B) the top 15 bacterial families (remaining families pooled as “Other”) for each carcass sample, faceted by carcass type and sampling day.

### Carcass type shapes microbial diversity and composition early in decomposition

Carcass type had its strongest influence early in decomposition. At Day 5, carcass type significantly predicted community composition (PERMANOVA, R^2^ = 0.099, p = 0.032) (Fig. 3), driven mainly by a mammal-reptile separation that did not survive multiple-testing correction (R^2^ = 0.112, p = 0.025, FDR-adjusted p = 0.075). By Day 9, carcass type no longer predicted composition (p = 0.223). Diversity followed a similar pattern: observed ASV richness differed significantly among carcass types at Day 5 (F = 3.82, df = 2,24, p = 0.036), with reptile carcasses supporting the most diverse communities, and Shannon diversity showed a similar but weaker trend (p = 0.075). These differences were accompanied by pronounced taxonomic differentiation among carcass types: 122 taxa were identified as indicators of a specific carcass type, and DESeq2 identified 25, 46, and 47 differentially abundant ASVs for the bird-mammal, bird-reptile, and mammal-reptile contrasts, respectively (Fig. 3E-G).

**Figure 3.**
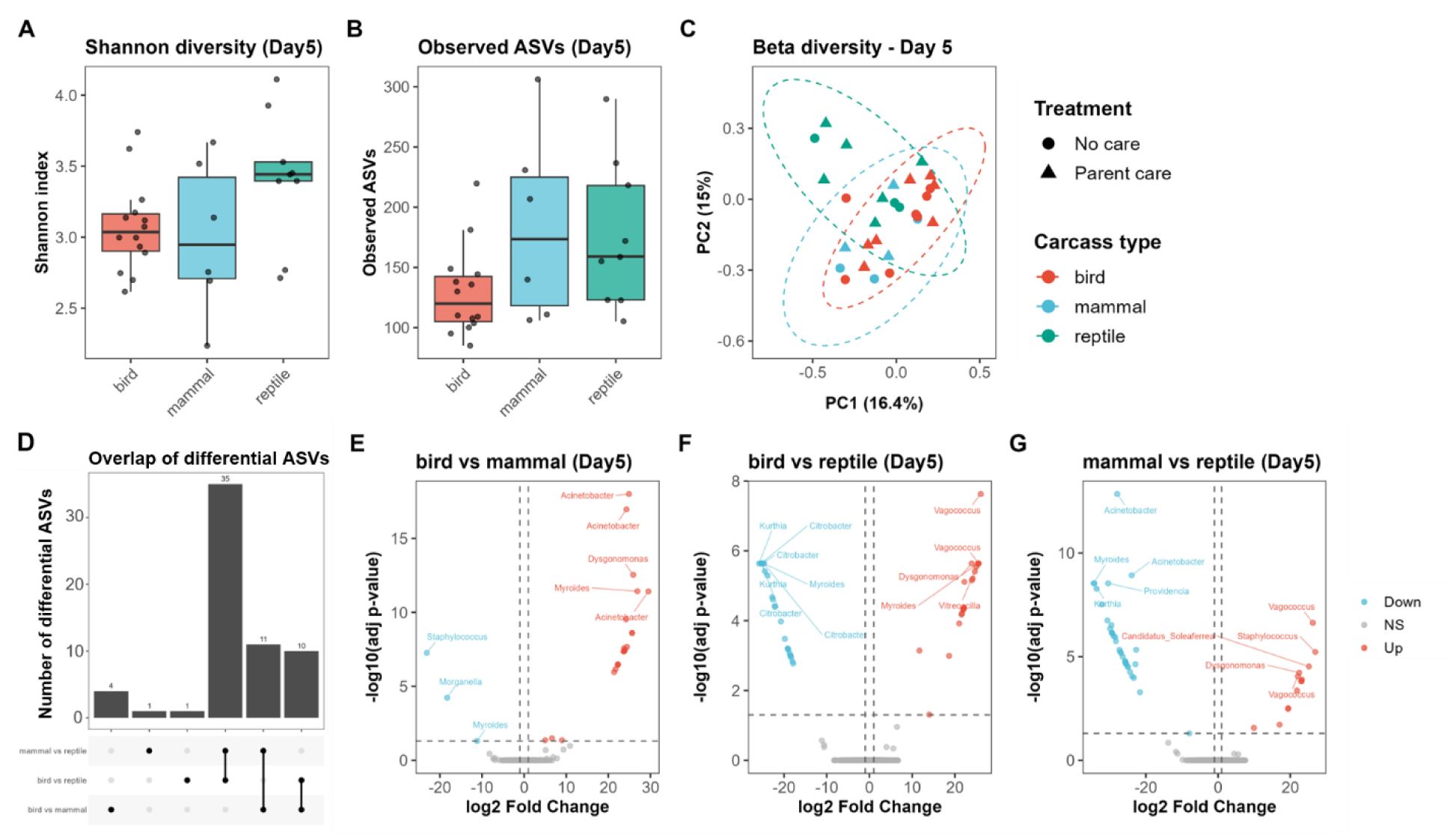
Carcass type shapes microbial diversity and composition at Day 5. (A) Shannon diversity, (B) observed ASV richness, (C) PCoA of Bray-Curtis dissimilarities at Day 5, (D) differentially abundant ASV overlap between carcass-type pairs, and (E-G) volcano plots of differentially abundant ASVs for each pairwise carcass-type comparison (bird vs. mammal, bird vs. reptile, mammal vs. reptile).

### Parental care effects emerge later in carcass-associated microbial communities

As carcass-type differences faded, a treatment signal began to emerge. By Day 9, treatment was significantly, though modestly, associated with carcass community composition once carcass type and batch were accounted for (PERMANOVA, R^2^ = 0.055, p = 0.045) (Fig. 4). Taxonomically, treatment differences were reflected by 58 differentially abundant ASVs and 18 indicator taxa (Fig. S4).

**Figure 4.**
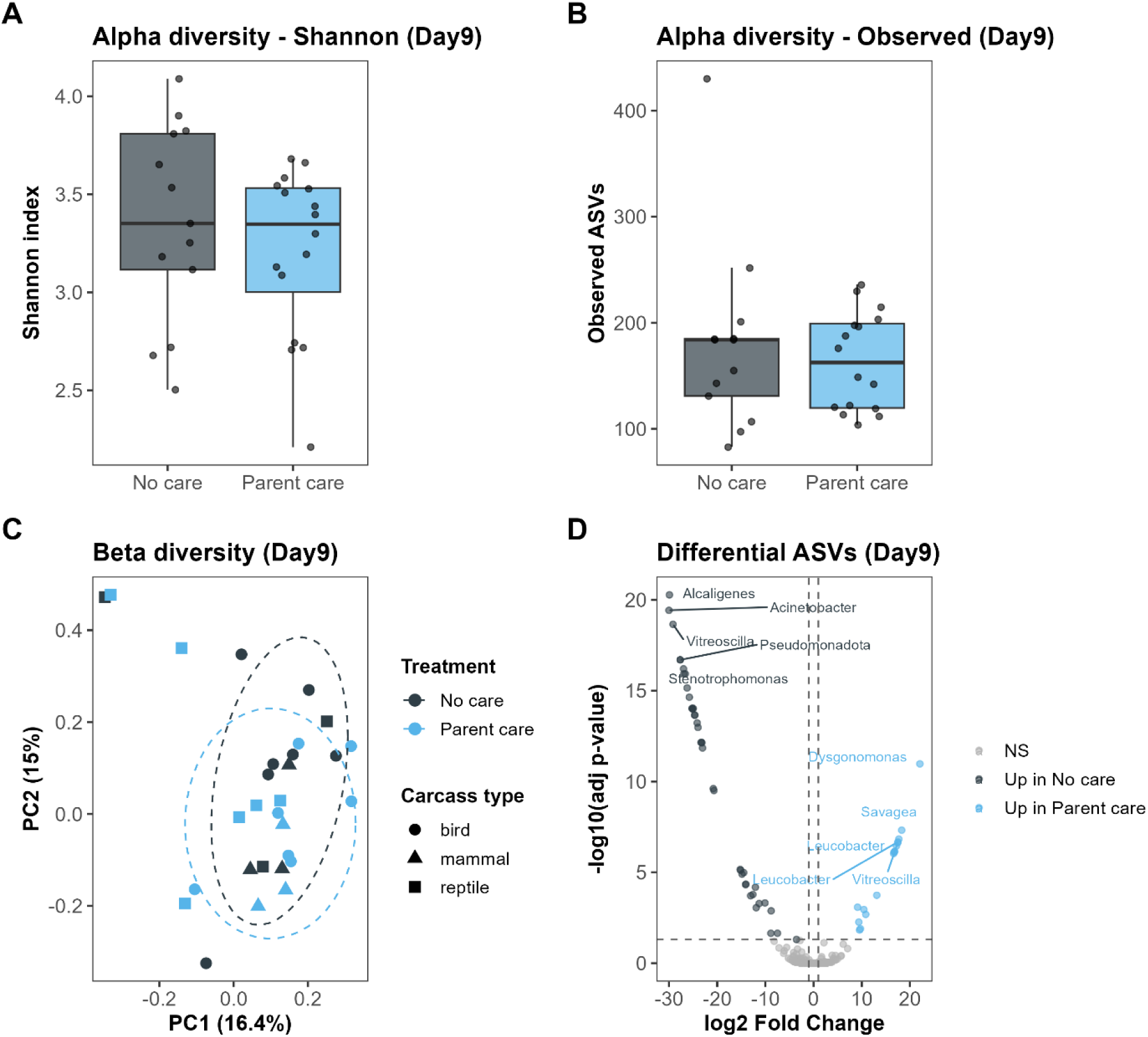
A modest treatment effect on carcass microbiome composition emerges by Day 9. (A) Shannon diversity, (B) observed ASV richness, (C) PCoA by treatment, and (D) volcano plot of differentially abundant ASVs. A heatmap of the top differentially abundant ASVs between treatments is shown separately in Supplementary Figure S4.

### The adult gut microbiome shows the strongest association with experimental treatment

This treatment signal was strongest not on the carcass itself, but in the adult beetle gut. The adult beetle gut microbiome was the most strongly treatment-structured compartment in the dataset (PERMANOVA, R^2^ = 0.281, p = 0.001) (Fig. S1), while carcass type had no detectable influence there (p = 0.324) and alpha diversity did not differ by treatment or carcass type. DESeq2 identified 74 differentially abundant ASVs between treatments (Fig. S5). Genus-level enrichment analysis revealed no significant enrichment after correcting for pseudoreplication (Fisher’s exact test, p = 0.167). Interpretation of this pattern is limited because No-care adults are necessarily sampled at Day 5 and Parent-care adults at Day 9, so treatment and sampling day are completely confounded in this compartment, and the effect cannot be attributed unambiguously to parental care itself as opposed to elapsed time or feeding duration.

### The larval gut microbiome shows little treatment or carcass-type signal

By contrast, the larval gut showed limited evidence of either signal, despite the strong reproductive effect of parental care reported above. Larval gut community composition was not significantly associated with treatment (p = 0.190) or carcass type (p = 0.127) (Fig. S2), with no clear biological structuring in this compartment beyond the sequencing-batch covariate (Table S1); these results were unchanged after excluding mammal carcasses. Shannon diversity differed among carcass types (p = 0.040) but not by treatment (p = 0.997).

### Cross-compartment and temporal linkages

Day 5 carcass and adult gut community structure were not correlated (partial Mantel r = -0.056, p = 0.704), whereas Day 5 carcass and larval gut structure showed a weaker but significant correlation (partial r = 0.212, p = 0.045), driven by Parent-care nests (partial r = 0.404, p = 0.006, n = 13; not assessable within No-care nests, n = 5). Adult and larval gut communities were not significantly correlated once treatment-group structure was accounted for (partial r = 0.232, p = 0.102) (Fig. 5); this result depended on the two available No-care nests, with a negative, non-significant correlation among the remaining Parent-care nests alone (r = -0.288, n = 8). Carcass structure at Day 5 remained correlated with Day 9 structure within the same nest, overall (partial r = 0.299, p = 0.004) and within Parent-care nests (partial r = 0.415, p = 0.002), but not within No-care nests alone (partial r = 0.188, p = 0.174). Within-nest microbiome turnover (Day 5 to Day 9 Bray-Curtis distance) did not differ between treatments (t-test, p = 0.118; Wilcoxon, p = 0.101) (Fig. S3).

**Figure 5.**
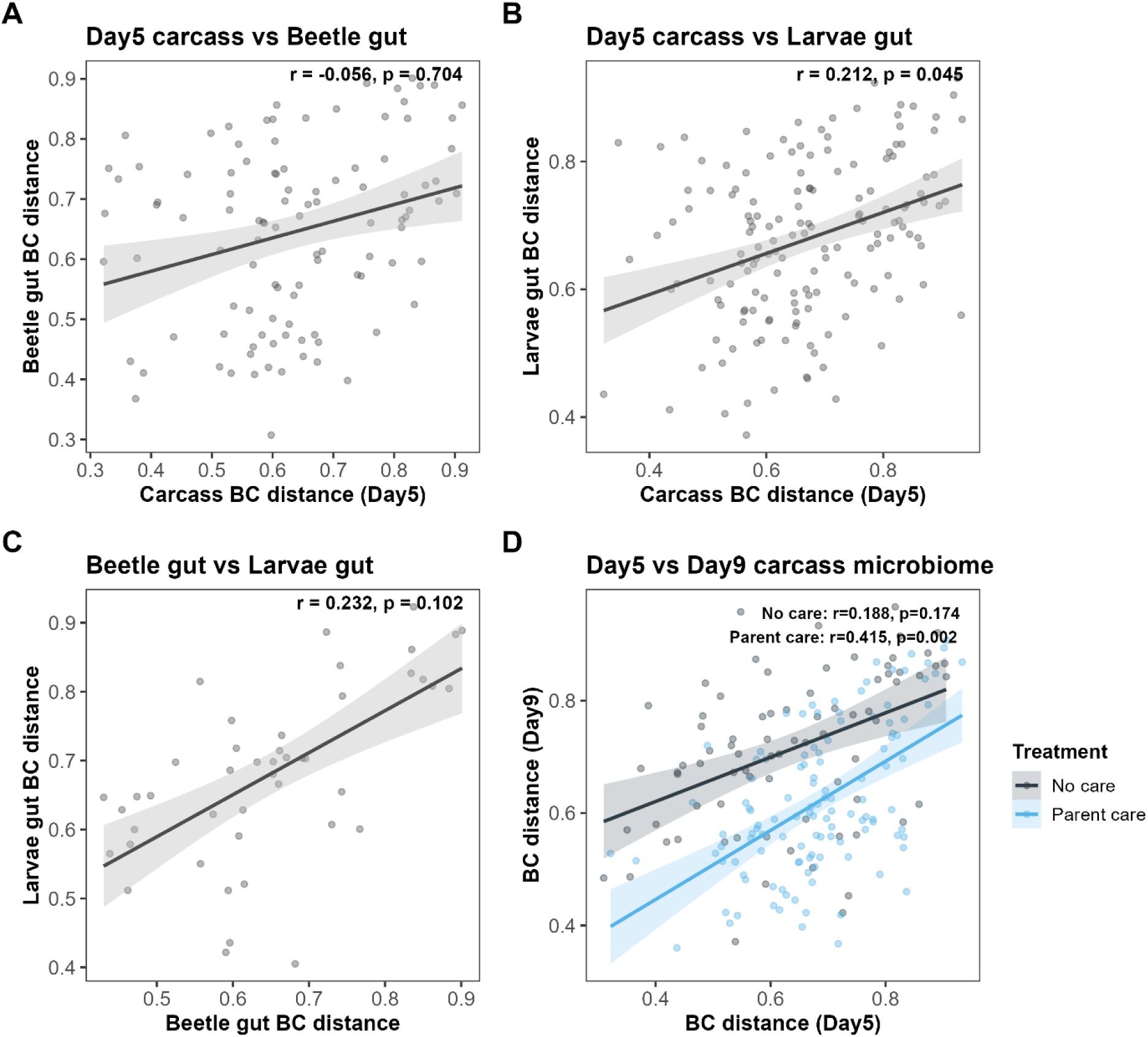
Cross-compartment and temporal linkages. Partial Mantel-test correlations for (A) Day 5 carcass vs. adult gut, (B) Day 5 carcass vs. larval gut, (C) adult gut vs. larval gut, and (D) Day 5 vs. Day 9 carcass. Regression lines show the unadjusted relationship between distance matrices for visualization.

## Discussion

### Main finding

Our study shows that carcass type and decomposition stage, rather than parental care per se, are the dominant drivers of carcass microbial community composition in this system, while parental care instead exerts its clearest and most robust effect on reproductive output itself. Rather than parental care progressively reshaping the carcass microbiome as decomposition proceeds, we find that carcass identity structures the microbiome from the outset, and that a treatment-by-success compositional interaction was not significant once carcass type and decomposition stage were accounted for. These results suggest that parental care may act primarily through adult-associated processes, most plausibly the mother’s own gut microbiome and behavior during care, rather than through large-scale restructuring of the carcass microbiome itself during early decomposition. Because microbial communities were sampled after treatment establishment, these associations should not be interpreted as causal effects. Breeding success, brood size, and brood mass were all robustly elevated by parental care across every sensitivity check we performed (e.g., adjusted OR = 16.1 for breeding success), whereas the corresponding treatment effect on carcass microbiome composition did not reach significance at Day 5 and remained modest (R² = 0.055) even by Day 9.

### Carcass identity and ecological realism

Carcass type was the strongest driver of microbial composition and diversity early in decomposition (Day 5), with reptile carcasses supporting a distinct, and generally more diverse, microbial community relative to bird and mammal carcasses. This pattern held even though all carcasses were frozen prior to use, suggesting it reflects underlying differences in carcass biochemistry across host taxonomic classes rather than a purely local/environmental artifact. To our best knowledge, this is among the first 16S rRNA datasets to characterize microbial succession on reptile carrion and to pair these data with the microbiome of a live burying beetle consumer. While *Nicrophorus* has occasionally been reported exploiting reptile resources such as snake eggs or reptile carcasses (Brown & Beresford 2016; Welch et al. 2023), these observations have not examined microbial community dynamics. Accordingly, our study extends *Nicrophorus* microbiome research beyond the laboratory vertebrate models that have dominated previous work.

This design also differs from most prior studies of parental care and the carcass microbiome, which have relied on a small number of standardized laboratory carcasses (e.g., Shukla et al 2018a, Shukla et al 2018b, Wang and Rozen 2017). Carcass identity was itself a strong predictor of both microbial composition and reproductive success in our dataset, indicating that the carcasses beetles breed on in nature present a much wider range of starting conditions than laboratory mice or chicks typically allow for. Despite this substantial natural heterogeneity, parental care consistently and robustly improved reproductive success, and this effect held up after excluding mammal carcasses, excluding laboratory-sourced carcasses, and under alternative modeling approaches (Supplementary Results). Parental care in this system therefore does not appear to depend on a standardized or homogeneous microbial starting point: females achieved consistently higher reproductive success across carcasses whose initial microbial communities differed substantially from one another.

This carcass-type effect extends beyond microbiome composition to reproductive success itself, apparently in tension with a previous analysis from this same laboratory of an overlapping wild-carcass panel, which concluded that carcass size, rather than source or taxonomic identity, determines breeding performance (Hsu et al 2024). We found that this apparent discrepancy is substantially explained by parental care: carcass type significantly predicted breeding success when parental care was experimentally withheld, but not when it was present, and a subset restricted to parental care and the original (pre-expansion) carcass batch reproduced a null carcass-type effect closely resembling that reported by Hsu et al. (2024). We interpret this as evidence that parental care buffers offspring against carcass-quality variation that is otherwise ecologically consequential in its absence, rather than as a genuine contradiction between the two studies; a formal carcass-type-by-parental-care interaction term did not reach significance in the pooled model, likely reflecting limited power in several sparsely sampled carcass-type-by-treatment cells, and we present the stratified comparison as the more informative test given this limitation.

### A compartment-specific microbiome response

By Day 9, a modest but significant treatment effect emerged on carcass composition, with the No-care community associated with genera including *Sphingobacterium*, *Myroides*, and *Alcaligenes*, and the Parent-care community associated with *Vagococcus* and *Tissierella* (Brunswick et al 2024). The adult beetle gut, however, showed by far the strongest treatment signal of any compartment sampled, with protective/antimicrobial genera such as *Carnobacterium*, *Pediococcus*, *Lactobacillus*, and *Enterococcus* more associated with No-care adults, and putrefactive genera including *Dysgonomonas*, *Ignatzschineria*, and *Morganella* more associated with Parent-care adults (Ammor et al 2006, Pechal et al 2013), though this enrichment pattern did not remain significant after correcting for pseudoreplication across ASVs of the same genus, and we report it as a descriptive, hypothesis-generating pattern. Because No-care adults are necessarily sampled at Day 5 and Parent-care adults at Day 9, treatment and sampling day are completely confounded in this compartment, and we cannot attribute this large effect unambiguously to parental care behavior itself as opposed to elapsed time or feeding duration; resolving this will require a design that samples adults from both treatments at a matched timepoint.

This treatment effect on the carcass reshaped which taxa were present without a corresponding change in overall community diversity (Table S1, S3). In marked contrast to its strong effect on reproduction, parental care left no detectable imprint on the larval gut: neither community composition (PERMANOVA, p = 0.190) nor Shannon diversity (p = 0.997) differed between care and no-care larvae, the latter an essentially complete null rather than a marginal non-significance. This is the compartment in which the extended-phenotype model most directly predicts an effect, because parents are proposed to seed offspring with a shared microbiota (Shukla et al. 2018b; Wang and Rozen 2017); its absence here, set against the large reproductive effect, indicates a microbially compartmentalized system rather than one in which care propagates a common microbiome to offspring. The only structuring signal detected in the larval gut was a carcass-type difference in Shannon diversity (p = 0.040) — higher in mammal- and reptile-than bird-associated larvae, though this reflects an omnibus test rather than formal pairwise contrasts. Whatever consequence parental care has for the microbiome in this system is therefore concentrated in the adult, not the offspring, compartment.

This absence of a detectable treatment effect in the larval gut appears, at first glance, to be in tension with Wang and Rozen (2017), who found that full- and prehatch-care larvae of *Nicrophorus vespilloides* were predominantly colonized by maternal gut bacteria, whereas no-care larvae instead acquired bacteria from the carcass. Two differences between the two studies may help explain this apparent discrepancy. First, the two studies examine different *Nicrophorus* species (*N. vespilloides* versus *N. nepalensis*), and species-specific differences in parental behavior or larval gut physiology could plausibly alter how strongly parental care shapes larval colonization. Second, Wang and Rozen (2017) tracked culturable bacterial isolates and asked whether individual strains recovered from larvae matched maternal- or carcass-derived isolates, whereas we characterized whole-community composition via 16S rRNA amplicon sequencing. A small number of readily culturable strains could transfer detectably from the mother to her offspring without producing a measurable shift in overall community composition, particularly if those strains are already present, even at low abundance, in both treatment groups’ background microbiota.

These two approaches are therefore not necessarily measuring the same thing, and our results do not rule out a component of maternal-strain transmission occurring alongside the compartmentalized, largely adult-localized pattern we report here.

### Cross-compartment isolation and the role of the attending female

Our compositional analyses reveal a highly compartmentalized microbial system. Contrary to the hypothesis that parents transmit a shared gut microbiota to their offspring, we found no robust cross-compartment correlations—such as between the adult and larval gut—once experimental treatments were properly accounted for. The only surviving linkage was a modest connection between the early-stage (Day 5) carcass and the larval gut, restricted entirely to parent-care nests. This compartmentalized pattern reframes the role of the attending female: rather than acting as a microbial conduit, she functions as an intermediary. The substantial microbial shifts occur primarily within the adult female’s gut, while the larval gut remains largely unaffected by treatment or carcass type. Thus, the female may represent a major site of microbial turnover rather than a stable donor, absorbing the compositional consequences of prolonged carcass management rather than transmitting a consistent microbiome onward. Under this view, parental care is fundamentally a mechanism of ecological carcass management, with the female’s microbiome shifting as a by-product of this intensive exposure.

### Temporal stability driven by parental care

Temporally, parental care alters the carcass microbiome not by drastically changing the magnitude of microbial turnover, but by stabilizing community identity over time. Within-nest microbiome turnover from Day 5 to Day 9 did not differ measurably between treatments. However, a nest’s Day 5 carcass community significantly predicted its Day 9 community only when parental care was present. In the absence of care, this early microbial signature fails to persist. This suggests that rather than completely overhauling the overarching decomposition process—which remains dominated by carcass type— parental care acts to stabilize the microbial environment. This temporal stabilization aligns well with established mechanisms of active carcass maintenance, such as the application of antimicrobial secretions, which effectively check unchecked microbial proliferation and maintain a predictable environment for the developing brood (Arce et al., 2013; Cotter et al., 2011; Duarte et al., 2018; Shukla et al., 2018a; Vogel et al., 2017).

### Limitations

We note four methodological considerations. First, to address the risk of quasi-complete separation caused by small sample sizes in certain carcass-by-treatment cells (e.g., n = 3 for successful parent-care mammals), we applied Firth’s penalized regression (Firth, 1993), though formal interaction tests may remain underpowered. Second, relying on opportunistic citizen-science donations of fresh, intact carcasses introduced sampling bias. Small mammals such as mice, shrews, and bats were rarely recovered in suitable condition compared to birds and reptiles, most likely due to earlier start of decaying process in abdomen and disarticulations in these taxa (e.g. Brand, Hussey and Taylor (2003)). In addition, small mammals recovered from glue traps were excluded because of possible poisoning or chemical contamination. Together, these factors explain their comparatively small and taxonomically narrow representation in our sample. Third, the uneven species count within vertebrate categories precluded modeling species identity as a random effect. We instead confirmed the robustness of carcass-category effects through sensitivity analyses that excluded mammalian and laboratory-sourced carcasses (Supplementary Methods). Fourth, the significant carcass-type PERMANOVA result (Table S1) was accompanied by a significant difference in within-group dispersion among carcass types (betadisper, p = 0.042; Table S8), indicating that this result partly reflects unequal within-group spread rather than centroid separation alone; treatment-related PERMANOVA results were not affected, as dispersion was homogeneous across treatment groups (Table S8).

## Conclusion

Together, these results indicate that the extended-phenotype consequences of parental care in this system operate less through a progressive, care-driven remodeling of the carcass microbiome, and more through a direct effect of care on reproductive output itself, alongside a comparatively modest and largely adult-localized microbiome signature that does not extend to the larval gut. A carcass-identity effect on both the microbiome and reproductive success persists alongside this direct effect and may be modulated, though not overridden, by parental care; disentangling the relative contributions of carcass identity, elapsed time, and parental behavior (ideally with a design that also varies carcass identity, timing of care removal, and sampling day independently) will be necessary to establish whether carcass microbiome state is a cause, a consequence, or simply a correlate of breeding success in this system.

Using carcasses spanning substantial natural microbial and taxonomic heterogeneity, our results suggest that parental care in burying beetles functions primarily as a direct driver of reproductive success and, to a lesser extent, of adult-gut microbiome composition, rather than as a mechanism that broadly restructures the carcass or larval microbiome.

## Acknowledgements

SJS was supported by Career Development Project (115L7845) and Seed Projects for Interdisciplinary Research (NTU-SPIR-115L8422) from National Taiwan University, National Science and Technology Council 2030 Cross-Generation Young Scholars Program (113-2628-B-002-028-; 114-2628-B-002-027-), and Yushan Fellow Program (MOE-111-YSFAG-0003-002-P1) provided by the Ministry of Education, Taiwan.

## Author contribution

WJL, CHH and SJS conceived the ideas;

WJL, CHH and SJS designed the experiments;

WJL, CHH and SJS collected the data;

WJL and SJS analyzed the data;

WJL and SJS wrote the first draft of the manuscript;

all authors revised the manuscript and approved the final version for publication.

## Data availability

The 16S rRNA amplicon sequencing data generated in this study will be deposited in a public repository (e.g., NCBI Sequence Read Archive) prior to publication; accession numbers will be provided upon deposition. All other data supporting the findings of this study are available within the paper and its Supplementary Information.

## Code availability

The R scripts used for statistical analyses in this study are available from the corresponding author upon reasonable request.

