## Supplemental data for "Parental care enhances reproductive success without remodeling the carcass microbiome in a wild burying beetle"

### Supplementary information

#### Methods

##### *Sensitivity and robustness analyses: reproductive outcomes*

Excluding the four laboratory-sourced carcasses (wild-caught only) did not change the direction or significance of the treatment or carcass-type effects on breeding success, indicating that carcass origin was not driving the reproductive results. Sequencing batch and log-transformed carcass mass were not significant predictors of success once treatment and carcass type were included. The mammal carcass-type effect on success, while significant, was substantially attenuated under Firth's penalized regression (OR = 26.8, further reduced to OR = 15.8 after excluding the two laboratory-mouse replicates) and in a bird-vs-reptile-only model (reptile OR = 13.5, p = 0.032). A simpler paired comparison restricted to same-species carcass pairs (Wilcoxon signed-rank) showed the same direction of effect as the covariate-adjusted models but did not reach significance for brood size (p = 0.142) or brood mass (p = 0.069), reflecting the lower power of a test that does not adjust for carcass type, batch, or carcass mass.

The treatment-dependent carcass-type effect on success (see Results) did not appear to be an artifact of reduced statistical power in the smaller original (batch 1 only) sample: restricting to batch 1 alone, carcass type remained a significant predictor of success with treatments pooled (Firth likelihood-ratio test, p = 0.005). It also did not reflect a nonlinear treatment of carcass mass: carcass type remained significant (p = 0.008) after adding a quadratic carcass-mass term, which itself significantly improved model fit (AIC 77.1 vs. 80.3 for the linear-only model). The formal carcass x treatment interaction term was not significant for either brood size (p = 0.797) or brood mass (p = 0.400).

Carcass mass, which spanned a 25-fold range (3.1-78.4 g), was log-transformed and added as a covariate to the logistic and linear models of reproductive outcome; its association with brood size and brood mass was additionally assessed with Spearman correlation to evaluate whether carcass size confounded the treatment and carcass-type effects.

##### *Sensitivity and robustness analyses: community composition and diversity*

Because the mammal category comprised only three species (n = 6 carcasses) and remained comparatively small even after batch 2 was added, every PERMANOVA, alpha-diversity model, and reproductive-outcome model involving carcass type was repeated after excluding mammal carcasses, to assess whether apparent carcass-type effects were disproportionately driven by this small group.

##### *Collinearity and model diagnostics*

Variance inflation factors (car::vif(), computed on the model design matrix independently of the response variable) were calculated for every multi-predictor model used in the community- and reproduction-level analyses. Residual normality for every linear or mixed model was checked with the Shapiro-Wilk test; where residuals departed from normality, results were interpreted alongside the corresponding non-parametric (Wilcoxon, Kruskal-Wallis) tests.

#### Supplementary Results

**Sensitivity analyses: reproductive outcomes**

Excluding the four laboratory-sourced carcasses (wild-caught only) did not change the direction or significance of the treatment or carcass-type effects on breeding success, indicating that carcass origin was not driving the reproductive results. Sequencing batch and log-transformed carcass mass were not significant predictors of success once treatment and carcass type were included. The mammal carcass-type effect on success, while significant, was substantially attenuated under Firth's penalized regression (OR = 26.8, further reduced to OR = 15.8 after excluding the two laboratory-mouse replicates) and in a bird-vs-reptile-only model (reptile OR = 13.5, p = 0.032). A simpler paired comparison restricted to same-species carcass pairs (Wilcoxon signed-rank) showed the same direction of effect as the covariate-adjusted models but did not reach significance for brood size (p = 0.142) or brood mass (p = 0.069), reflecting the lower power of a test that does not adjust for carcass type, batch, or carcass mass.

The treatment-dependent carcass-type effect on success did not appear to be an artifact of reduced statistical power in the smaller original (batch 1 only) sample: restricting to batch 1 alone, carcass type remained a significant predictor of success with treatments pooled (Firth likelihood-ratio test, p = 0.005). It also did not reflect a nonlinear treatment of carcass mass: carcass type remained significant (p = 0.008) after adding a quadratic carcass-mass term, which itself significantly improved model fit (AIC 77.1 vs. 80.3 for the linear-only model). The formal carcass x treatment interaction term was not significant for either brood size (p = 0.797) or brood mass (p = 0.400).

**Sensitivity analyses: community composition and diversity**

Because the mammal category comprised only three species (n = 6 carcasses), every PERMANOVA and alpha-diversity model involving carcass type was repeated after excluding mammal carcasses. The Day 5 carcass-type effect on composition weakened to a trend when mammal carcasses were excluded (bird vs. reptile only, p = 0.063), indicating the full three-group result was partly, but not entirely, driven by the mammal group; the effect on richness strengthened after exclusion (F = 8.43, df = 1,19, p = 0.009), and the effect on Shannon diversity also strengthened (bird-vs-reptile only, p = 0.026). The Day 9 treatment effect on composition was reduced to a non-significant trend after excluding mammal carcasses (p = 0.107), but the effect size was essentially unchanged (R = 0.055 vs. 0.061), indicating that this reflects reduced statistical power rather than the effect being an artifact of the mammal group. The adult-gut treatment effect was essentially unchanged after excluding mammal carcasses (R = 0.286, p = 0.002). Larval-gut results were unchanged after excluding mammal carcasses.

#### *Partial Mantel tests for cross-compartment comparisons*

Day 5 carcass and adult-gut samples were often processed within the same sequencing batch, and batch was consistently among the strongest predictors of community composition throughout this dataset. Because a shared batch could inflate the apparent similarity between compartments independent of any genuine biological association, cross-compartment Mantel comparisons (Day 5 carcass vs. adult gut, Day 5 carcass vs. larval gut, adult gut vs. larval gut) were tested as partial Mantel tests (vegan::mantel.partial(), 999 permutations), conditioning on a batch-difference matrix (0 for sample pairs from the same sequencing batch, 1 otherwise). The batch-controlled statistic is reported throughout the main text and Figure 5 as the primary cross-compartment result, except for the adult gut-larval gut comparison. Because this correlation remained significant after conditioning on batch alone, and because the sample available for this comparison was small and treatment-imbalanced (2 No-care, 8 Parent-care nests) -- raising the possibility that a spurious correlation could arise if the two treatment groups simply form distinct clusters in both compartments, independent of any genuine nest-level correspondence -- we additionally conditioned on a treatment-difference matrix (0 for sample pairs from the same treatment, 1 otherwise) and performed a leave-one-out analysis, removing each nest from the comparison in turn. The correlation fell to a non-significant level once treatment was conditioned upon (partial r = 0.232, p = 0.102), and the leave-one-out analysis showed the batch-only estimate depended on the two No-care nests: removing both left a negative, non-significant correlation among the remaining Parent-care nests alone (r = -0.288, n = 8). We therefore report the treatment-controlled statistic, rather than the batch-only statistic, as the primary result for the adult gut-larval gut comparison in the main text, Figure 5, and Table S7.

**Investigation of the sequencing-batch effect**

Because the second sequencing batch consisted almost entirely of reptile carcasses, we investigated whether the batch effect on community composition was a genuine technical signal or an artifact of carcass-type composition. Homogeneity of multivariate dispersion did not differ between batches in any subset (betadisper, all p > 0.59), ruling out a dispersion-only artifact. Restricting the analysis to reptile carcasses only (holding carcass type constant), batch remained a significant predictor of composition (combined Day 5 + Day 9, R = 0.219, p = 0.001; Day 5 only, R = 0.316, p = 0.018; Day 9 only, R = 0.245, p = 0.070, likely underpowered at n = 9 per batch), supporting a genuine technical batch effect independent of carcass type.

**Model diagnostics**

Variance inflation factors for every multi-predictor model were below 2 in all cases, indicating that the modest collinearity between treatment and success reflects shared biological/statistical variance in the response rather than a design-matrix collinearity problem. Residual normality (Shapiro-Wilk) was acceptable for most linear models ($p > 0.10$), with two exceptions -- the Day 9 observed-richness model and the carcass-mass-adjusted brood-size and brood-mass models -- for which the corresponding non-parametric tests (Kruskal-Wallis, Wilcoxon) gave consistent conclusions.

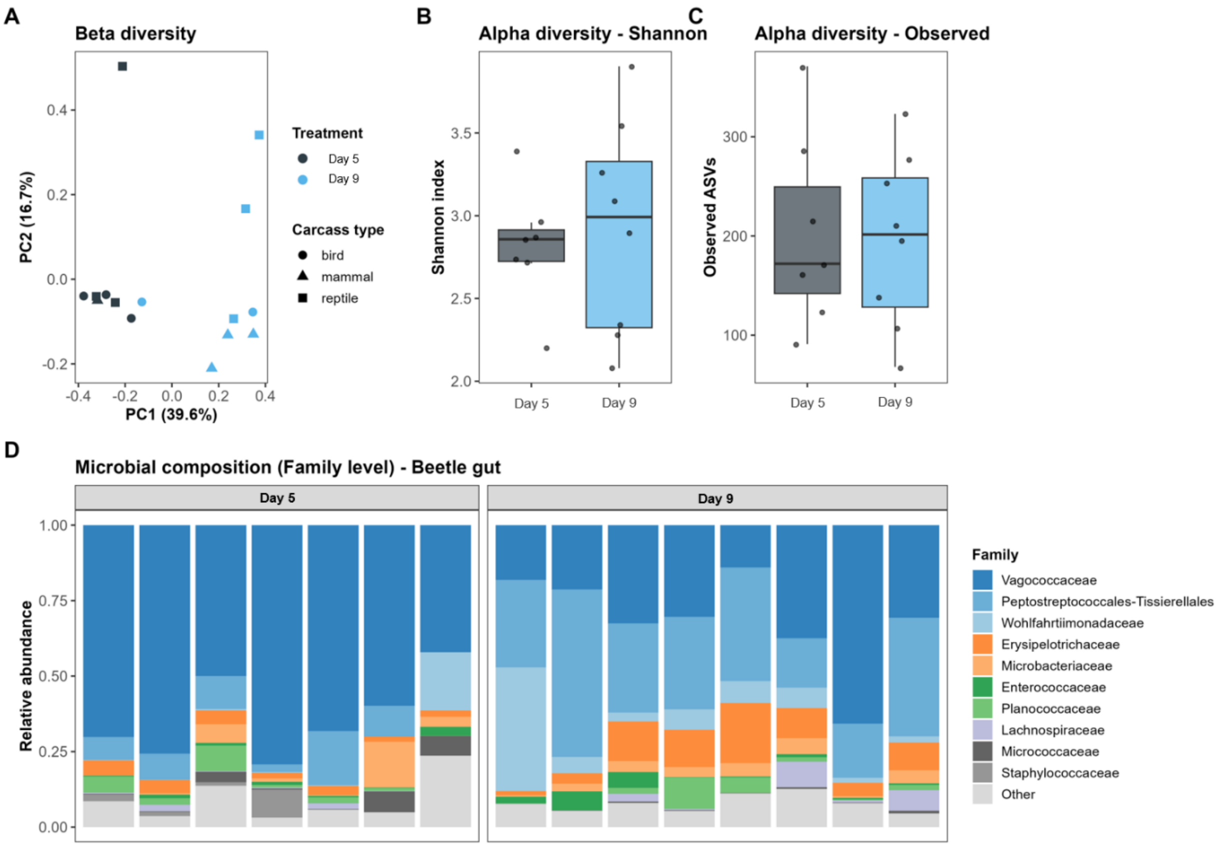


**Supplementary Figure S1**

Adult beetle gut microbiome. (A) PCoA of Bray-Curtis dissimilarities among adult gut communities, colored by day and shaped by carcass type. (B) Shannon diversity index and (C) observed ASV richness by day. (D) Relative abundance of bacterial families in the adult gut, by day.

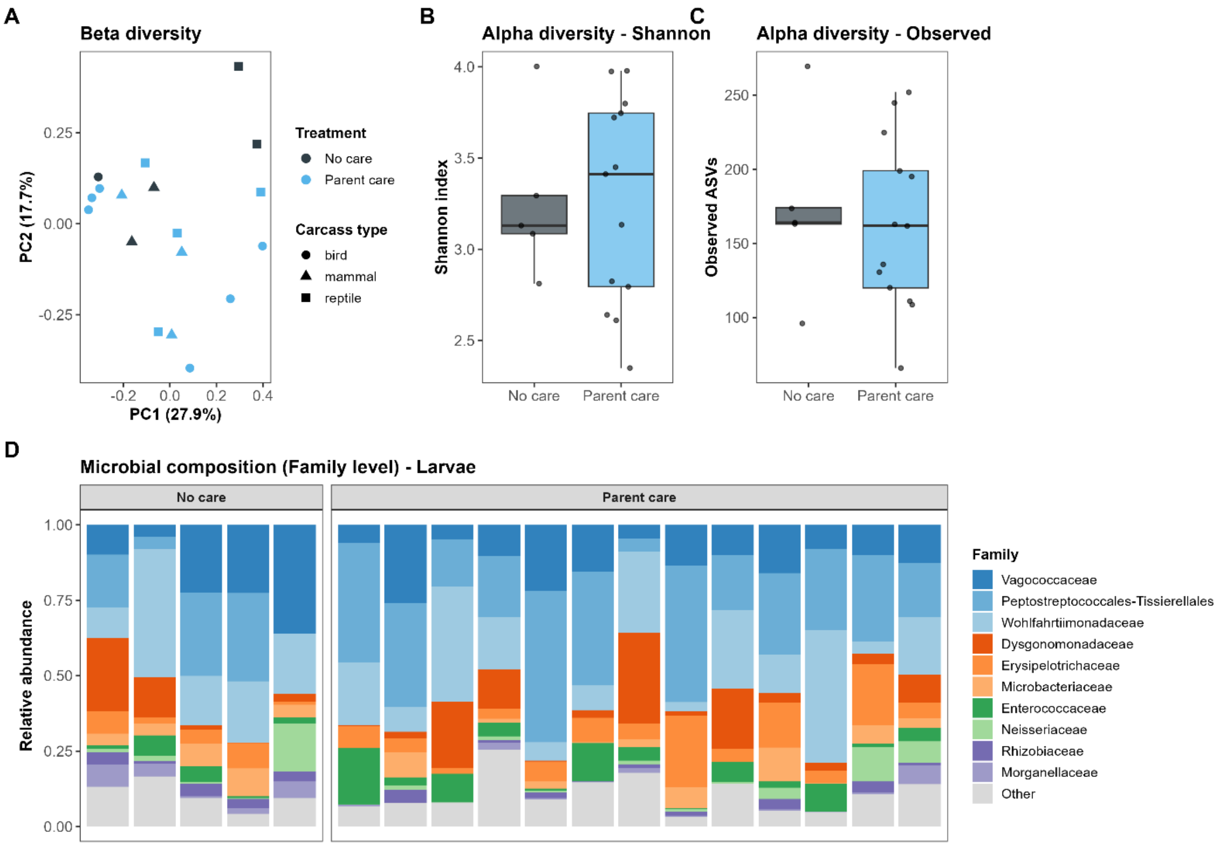


**Supplementary Figure S2**

Larval gut microbiome. (A) PCoA of Bray-Curtis dissimilarities among larval gut communities, colored by treatment and shaped by carcass type. (B) Shannon diversity index and (C) observed ASV richness by treatment. (D) Relative abundance of bacterial families in the larval gut, by treatment.

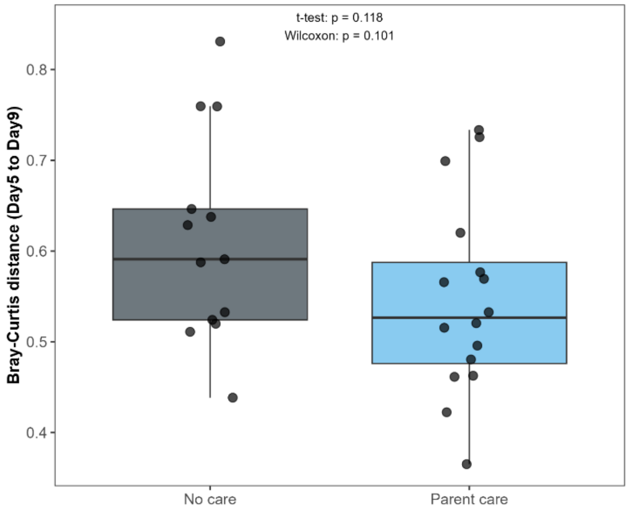


**Supplementary Figure S3**

Within-nest carcass microbiome turnover from Day 5 to Day 9. Bray-Curtis distance between the Day 5 and Day 9 carcass community of the same nest, compared between the No care and Parent care treatments.

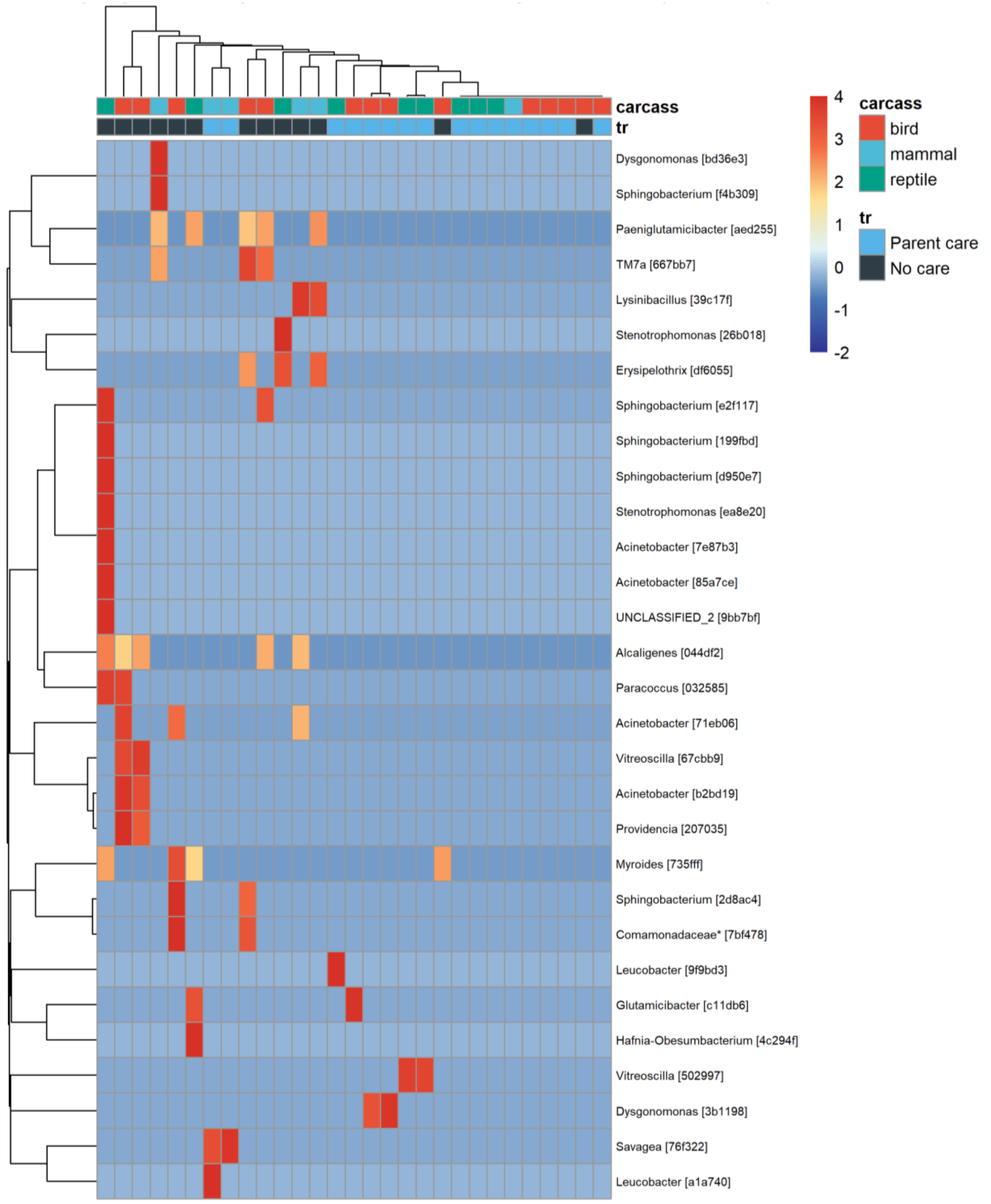


**Supplementary Figure S4**

Differentially abundant ASVs between treatments at Day 9. Heatmap of the top 30 ASVs ranked by adjusted p-value (DESeq2 Wald test), with samples (columns) annotated by carcass type and treatment; color indicates row-scaled (z-score) relative abundance. Taxon labels reflect the finest classification resolved for each ASV: unmarked labels are genus-level assignments (shown in italics, including provisionally named candidate genera such as TM7a); * denote family- labels used instead when genus-level classification was not available. This convention applies to all heatmaps in the main text and Supplementary Figures.

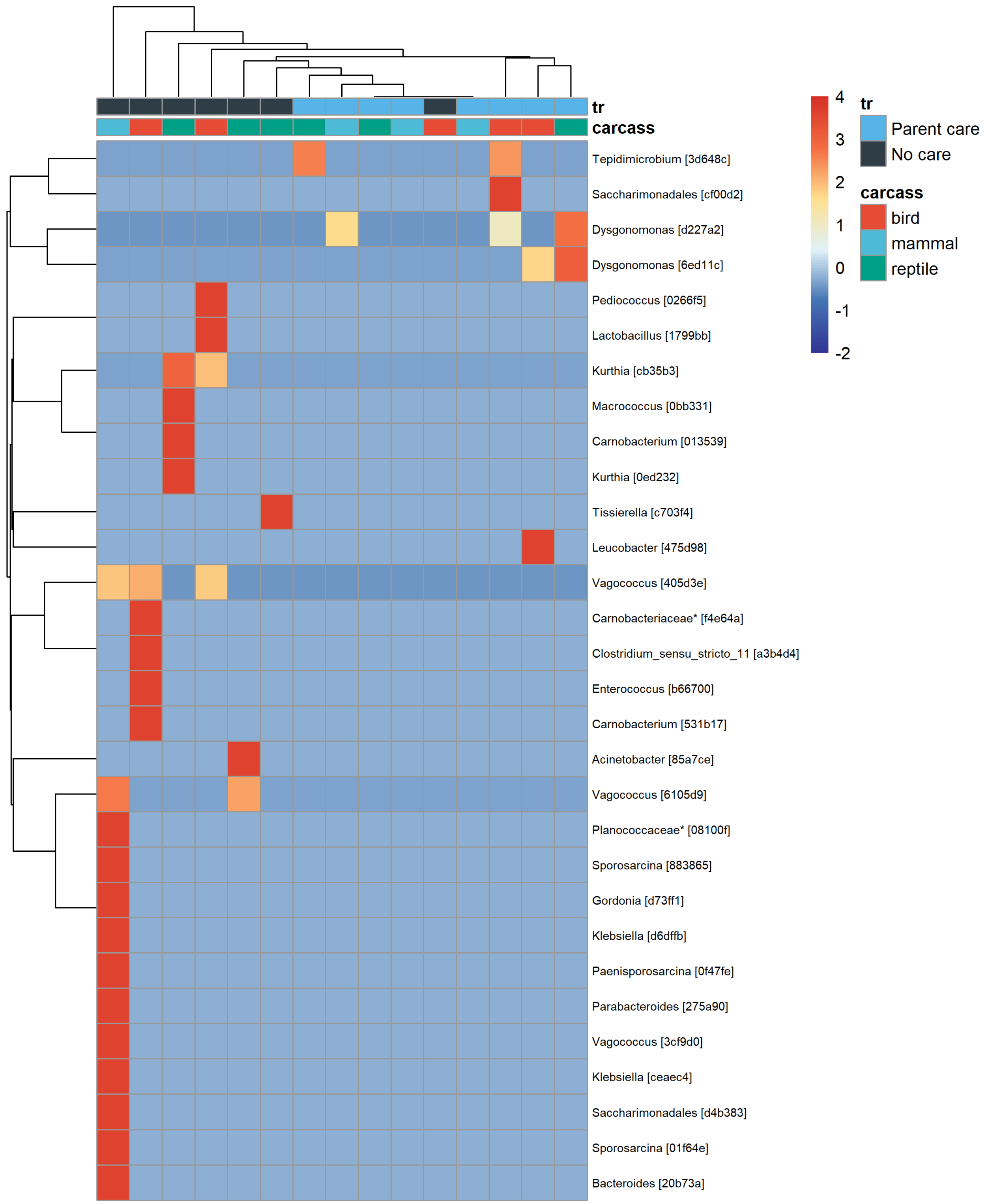


**Supplementary Figure S5**

Differentially abundant ASVs between treatments in the adult beetle gut. Heatmap of the top 30 ASVs ranked by adjusted p-value (DESeq2 Wald test), with samples (columns) annotated by carcass type and treatment; color indicates row-scaled (z-score) relative abundance; * denote family- labels used instead when genus-level classification was not available. This convention applies to all heatmaps in the main text and Supplementary Figures.

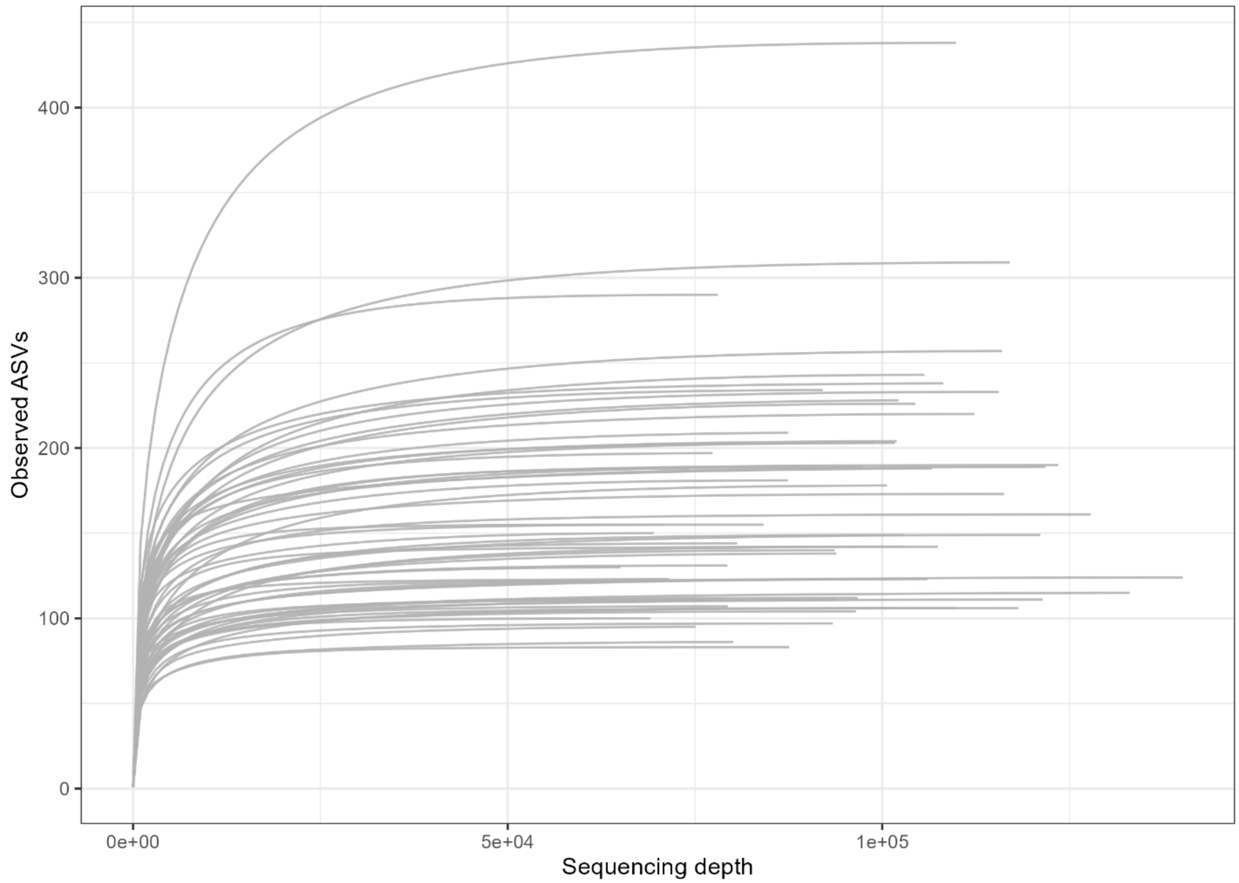


**Supplementary Figure S6**

Rarefaction curves. Observed ASV richness as a function of sequencing depth for every sample across all compartments (carcass, adult gut, larval gut), confirming that sequencing depth was sufficient to approach saturation of richness prior to rarefaction.

### Supplementary Tables

**Table S1. PERMANOVA (ADONIS) results for microbial community composition -- full model output**

| Dataset / model | Term | Df | R² | F | p value |
|---|---|---|---|---|---|
| Full dataset (carcass+tr+group+batch) | Carcass type | 2 | 0.056 | 2.04 | **0.003** |
| Full dataset (carcass+tr+group+batch) | Treatment | 1 | 0.018 | 1.32 | 0.168 |
| Full dataset (carcass+tr+group+batch) | Decomposition stage (group) | 1 | 0.093 | 6.73 | **0.001** |
| Full dataset (carcass+tr+group+batch) | Sequencing batch | 1 | 0.070 | 5.07 | **0.001** |
| Day 5 (carcass+batch) | Carcass type | 2 | 0.099 | 1.66 | **0.032** |
| Day 5 (carcass+batch) | Sequencing batch | 1 | 0.100 | 3.36 | **0.001** |
| Day 5, success model (success+carcass+batch) | Success | 1 | 0.024 | 0.79 | 0.709 |
| Day 5, success model (success+carcass+batch) | Carcass type | 2 | 0.094 | 1.57 | 0.053 |
| Day 5, success model (success+carcass+batch) | Sequencing batch | 1 | 0.100 | 3.31 | **0.001** |
| Day 9 (carcass+tr+batch) | Carcass type | 2 | 0.075 | 1.20 | 0.223 |
| Day 9 (carcass+tr+batch) | Treatment | 1 | 0.054 | 1.75 | **0.045** |
| Day 9 (carcass+tr+batch) | Sequencing batch | 1 | 0.081 | 2.61 | **0.008** |
| Day 9, success model (success+tr+carcass+batch) | Success | 1 | 0.041 | 1.35 | 0.176 |
| Day 9, success model (success+tr+carcass+batch) | Treatment | 1 | 0.045 | 1.46 | 0.119 |
| Day 9, success model (success+tr+carcass+batch) | Carcass type | 2 | 0.069 | 1.12 | 0.302 |
| Day 9, success model (success+tr+carcass+batch) | Sequencing batch | 1 | 0.081 | 2.64 | **0.006** |
| Full dataset, tr x success interaction model | Sequencing batch | 1 | 0.107 | 6.90 | **0.001** |
| Full dataset, tr x success interaction model | Treatment x Success | 1 | 0.017 | 1.08 | 0.346 |
| Day 9, carcass x treatment interaction model | Sequencing batch | 1 | 0.088 | 2.79 | **0.007** |
| Day 9, carcass x treatment interaction model | Carcass type x Treatment | 2 | 0.048 | 0.76 | 0.801 |
| Adult beetle gut (tr+carcass+batch) | Treatment | 1 | 0.281 | 6.87 | **0.001** |
| Adult beetle gut (tr+carcass+batch) | Carcass type | 2 | 0.093 | 1.13 | 0.324 |
| Adult beetle gut (tr+carcass+batch) | Sequencing batch | 1 | 0.115 | 2.81 | **0.020** |
| Adult beetle gut, carcass x treatment interaction model | Sequencing batch | 1 | 0.115 | 2.86 | **0.013** |
| Adult beetle gut, carcass x treatment interaction model | Carcass type x Treatment | 2 | 0.089 | 1.12 | 0.342 |
| Larval gut (tr+carcass+batch) | Treatment | 1 | 0.069 | 1.31 | 0.190 |
| Larval gut (tr+carcass+batch) | Carcass type | 2 | 0.141 | 1.35 | 0.127 |
| Larval gut (tr+carcass+batch) | Sequencing batch | 1 | 0.101 | 1.93 | **0.039** |

*PERMANOVA (adonis2, 999 permutations, marginal/Type-III term testing) on Bray-Curtis dissimilarities, transcribed directly from the adonis2() output in the analysis console log (each block = one fitted model; rows below the same model label are terms from that same model, tested jointly). Significant terms ($p \leq 0.05$) are in bold.*

**Table S2. Pairwise PERMANOVA comparisons of carcass type and treatment**

| Comparison | $R^2$ | p value | p (FDR-adjusted) |
|---|---|---|---|
| Day 5 carcass type: bird vs. mammal | 0.077 | 0.155 | 0.155 |
| Day 5 carcass type: bird vs. reptile | 0.062 | 0.063 | 0.095 |
| Day 5 carcass type: mammal vs. reptile | 0.112 | **0.025** | 0.075 |
| Day 9 treatment: No care vs. Parent care | 0.055 | 0.052 | 0.052 |

*Pairwise carcass-type/treatment comparisons use false-discovery-rate (Benjamini-Hochberg) correction across the 3 (Day5) or single (Day9) comparisons within each set. Significant terms ($p \leq 0.05$) are in bold. For the Day 9 treatment row, this test controls for sequencing batch but, unlike the marginal model underlying Table S1, does not additionally control for carcass type; the Table S1 model, which does control for carcass type, gave a nearly identical result for the treatment term ($R^2 = 0.054$, $p = 0.045$).*

**Table S3. Effects of carcass type, treatment, and batch on alpha diversity (Shannon index, observed ASV richness)**

| Compartment | Response | Term | Df | F | p value |
|---|---|---|---|---|---|
| Day 5 carcass | Shannon | Carcass type | 2 | 2.89 | 0.075 |
| Day 5 carcass | Shannon | Treatment | 1 | 0.34 | 0.567 |
| Day 5 carcass | Shannon | Sequencing batch | 1 | 6.28 | **0.019** |
| Day 5 carcass | Observed ASVs | Carcass type | 2 | 3.82 | **0.036** |
| Day 5 carcass | Observed ASVs | Treatment | 1 | 0.02 | 0.882 |
| Day 5 carcass | Observed ASVs | Sequencing batch | 1 | 9.12 | **0.006** |
| Day 9 carcass | Shannon | Carcass type | 2 | 0.76 | 0.477 |
| Day 9 carcass | Shannon | Treatment | 1 | 0.51 | 0.483 |
| Day 9 carcass | Shannon | Sequencing batch | 1 | 2.05 | 0.165 |
| Day 9 carcass | Observed ASVs | Carcass type | 2 | 0.91 | 0.416 |
| Day 9 carcass | Observed ASVs | Treatment | 1 | 0.23 | 0.635 |
| Day 9 carcass | Observed ASVs | Sequencing batch | 1 | 0.77 | 0.389 |
| Adult beetle gut | Shannon | Treatment | 1 | 0.22 | 0.650 |
| Adult beetle gut | Shannon | Carcass type | 2 | 2.13 | 0.170 |
| Adult beetle gut | Shannon | Sequencing batch | 1 | 5.31 | **0.044** |
| Larval gut | Shannon | Treatment | 1 | 0.00 | 0.997 |
| Larval gut | Shannon | Carcass type | 2 | 4.18 | **0.040** |
| Larval gut | Shannon | Sequencing batch | 1 | 9.30 | **0.009** |

*Linear models on rarefied alpha-diversity metrics (ANOVA Type-I sums of squares in the order carcass/treatment/batch shown; term order does not change significance here as designs are balanced-ish). Sensitivity check: excluding the 3 mammal-carcass species from the Day 5 model does not weaken these results -- Shannon carcass F = 5.80, p = 0.026; Observed ASVs carcass F = 8.43, p = 0.009 -- making the Day-5 carcass-type effect on diversity the most robust result to sample-size perturbation in the dataset. Significant terms (p ≤ 0.05) are in bold.*

**Table S4. Effects of parental care, carcass type, and batch on reproductive outcomes**

| Response variable | Model / test | Statistic | p value |
|---|---|---|---|
| Breeding success | Fisher's exact test (treatment) | OR = 8.8 | **< 0.0001** |
| Breeding success | Binomial GLM, treatment (adj. carcass type + batch) | OR = 16.1 (95% CI 4.6-77.9) | **< 0.0001** |
| Breeding success | Binomial GLM, carcass type: mammal vs. bird | OR = 44.2 (95% CI 3.9-1205) | **0.006** |
| Breeding success | Binomial GLM, carcass type: reptile vs. bird | OR = 13.8 (95% CI 1.4-184) | **0.031** |
| Breeding success | Binomial GLM, sequencing batch | OR = 0.18 (95% CI 0.015-1.55) | 0.132 |
| Breeding success | Family GLMM (crossed random effects), treatment | chi-sq = 15.5, df = 1 | **< 0.0001** |
| Breeding success | Family GLMM, carcass type (Type II Wald) | chi-sq = 9.65, df = 2 | **0.008** |
| Breeding success | Family GLMM, sequencing batch (Type II Wald) | chi-sq = 2.27, df = 1 | 0.132 |
| Brood size | Linear model (treatment, adj. carcass type + batch) | F = 15.6, df = 1,70 | **< 0.001** |
| Brood size | Family GLMM (Poisson), treatment (Type II Wald) | chi-sq = 132.4, df = 1 | **< 2e-16** |
| Brood size | Family GLMM (Poisson), carcass type (Type II Wald) | chi-sq = 6.94, df = 2 | **0.031** |
| Brood size | Family GLMM (Poisson), sequencing batch (Type II Wald) | chi-sq = 1.30, df = 1 | 0.254 |
| Brood mass | Linear model (treatment, adj. carcass type + batch + carcass mass) | F = 19.8, df = 1,70 | **< 0.001** |
| Brood mass | Linear model, carcass type (secondary predictor) | - | **0.015-0.036** |
| Brood mass | Family GLMM (lmer, incl. failed/zero broods), treatment (Type II Wald) | chi-sq = 21.3, df = 1 | **< 0.0001** |
| Brood mass | Family GLMM, successful broods only, treatment (Type II Wald) | chi-sq = 1.61, df = 1 | 0.204 |
| Mean larval mass | Treatment | - | 0.29 |
| Mean larval mass | Carcass mass (log) | - | **0.024** |

*The family-GLMM rows are a robustness check mirroring the original submission's random-effects design (crossed male/female family identity); the simple treatment/carcass/batch linear models above them are the primary reported models. The "successful broods only" row for brood mass shows that once a nest succeeds, treatment no longer predicts how large the brood mass is (chi-sq = 1.61, p = 0.204). Significant terms (p ≤ 0.05) are in bold.*

**Table S5. Carcass type x treatment interaction on breeding success**

| Comparison | Test | Statistic | p value |
|---|---|---|---|
| Pooled model, carcass type x treatment | Likelihood-ratio test | - | 0.827 |
| No-care group, carcass type effect | Firth penalized-regression LRT | - | **0.018** |
| Mammal vs. bird (No-care) | Binomial GLM | OR = 26.0 | **0.042** |
| Reptile vs. bird (No-care) | Binomial GLM | OR = 26.0 | **0.042** |
| Parent-care group, carcass type effect | Firth penalized-regression LRT | - | 0.248 |
| Original batch (batch1) only (both treatments), carcass type | Firth penalized-regression LRT | - | **0.005** |
| Full dataset + quadratic carcass-mass term | Carcass type retained | - | **0.008** |

*Carcass type was re-examined separately within the No-care and Parent-care subsets because several carcass-by-treatment cells were small enough to risk quasi-complete separation in standard logistic regression; Firth's penalized regression was used throughout. Significant terms ($p \leq 0.05$) are in bold.*

**Table S6. Differential abundance (DESeq2) and indicator species summary**

| Compartment / contrast | DESeq2 differentially abundant ASVs | Indicator taxa (multipatt) |
|---|---|---|
| Day 5, bird vs. mammal | 25 | - |
| Day 5, bird vs. reptile | 46 | - |
| Day 5, mammal vs. reptile | 47 | - |
| Day 5, any carcass type (total) | - | 122 |
| Day 9, treatment | 58 | 18 (14 No-care, 4 Parent-care) |
| Day 9, breeding success (fail vs. success) | - | 15 (7 fail-associated, 8 success-associated) |
| Adult beetle gut, treatment | 74 | 12 (3 No-care, 9 Parent-care) |

*Differential abundance was tested with DESeq2 (Benjamini-Hochberg adjusted $p < 0.05$, design including batch as a covariate); indicator taxa were identified with the IndVal.g statistic (indicspecies::multipatt(), 999 permutations, filtered to $p < 0.05$).*

**Table S7. Cross-compartment and temporal Mantel test correlations, and Day5-to-Day9 turnover**

| Comparison | Partial Mantel r | p value |
|---|---|---|
| Day 5 carcass vs. adult beetle gut | -0.056 | 0.704 |
| Day 5 carcass vs. larval gut | 0.212 | **0.045** |
| Day 5 carcass vs. larval gut, Parent-care only | 0.404 | **0.006** |
| Day 5 carcass vs. larval gut, No-care only (n=5) | -0.237 | 0.825 |
| Adult beetle gut vs. larval gut (batch only) | 0.557 | **0.012** |
| Adult beetle gut vs. larval gut (batch + treatment) | 0.232 | 0.102 |
| Day 5 vs. Day 9 carcass (all nests) | 0.299 | **0.004** |
| Day 5 vs. Day 9 carcass, No-care nests only | 0.188 | 0.174 |
| Day 5 vs. Day 9 carcass, Parent-care nests only | 0.415 | **0.002** |

*Partial Mantel tests (999 permutations), controlling for sequencing batch (and, for the adult beetle gut vs. larval gut comparison, additionally for treatment structure; see Supplementary Methods), on Bray-Curtis dissimilarities. The correlation between Day 5 carcass and adult beetle gut communities was not significant once batch was accounted for. The correlation between Day 5 carcass and larval gut communities was weaker but remained significant overall, and was significant when restricted to Parent-care nests; it could not be reliably assessed within No-care nests (n = 5). Turnover magnitude itself (paired Bray-Curtis distance, Day5 to Day9) did not differ by treatment: No-care mean = 0.613 (SD 0.115, n = 13) vs. Parent-care mean = 0.547 (SD 0.106, n = 16); t-test p = 0.118, Wilcoxon p = 0.101, linear model adjusting for carcass type and batch p = 0.158. Significant correlations (p <= 0.05) are in bold.*

**Table S8. Homogeneity of multivariate dispersion (betadisper), supporting the PERMANOVA analyses**

| Dataset | Grouping factor | Df | F | p value |
|---|---|---|---|---|
| All samples (Day5+Day9) | Carcass type | 2 | 3.36 | **0.042** |
| All samples (Day5+Day9) | Treatment | 1 | 0.26 | 0.610 |
| Day 5 carcass | Carcass type | 2 | 2.07 | 0.140 |
| Day 9 carcass | Treatment | 1 | 1.89 | 0.170 |
| Adult beetle gut | Treatment | 1 | 1.00 | 0.366 |
| Larval gut | Treatment | 1 | 0.52 | 0.492 |

*permutest(betadisper(), 999 permutations) tests whether groups differ in within-group dispersion rather than centroid location; PERMANOVA can conflate the two. Only one test is significant: carcass type across all carcass samples (p = 0.042), meaning the carcass-type PERMANOVA result (Table S1) partly reflects unequal within-group spread rather than centroid separation alone; treatment dispersion was homogeneous throughout, so the treatment PERMANOVA results are not confounded by this issue.*
